# DrosoTracker: a web application with a self-calibrating thermal model for husbandry scheduling and lifespan analysis in *Drosophila melanogaster*

**DOI:** 10.64898/2026.08.10.743933

**Authors:** Gerson Smith Asti Tello, Mariana Melani, Ana Clara Liberman

## Abstract

Planning husbandry tasks and experiments with Drosophila melanogaster requires converting a target date into development times that depend on the rearing temperature. This calculation needs to be done for each cross, genotype, and temperature, and the risk of error grows quickly. Available laboratory management tools let users register stocks, crosses, and track them, but they do not create schedules based on a clear, adjustable thermal model. To fill that gap, we developed DrosoTracker, a self-contained web application that works offline and predicts Drosophila development with a thermal summation model recalibrated through regression on data from Powsner (1935) (T0 = 11.78 °C, DD = 116.38 °C·days, R² = 0.997). The model offers an optional two-level calibration driven by user observations. A wild-type strain first adjusts the model to the laboratory’s own conditions. Then each genotype is calibrated against that reference using a random-effects shrinkage estimator that accounts for measurement error and between-batch variability. The model creates schedules for husbandry tasks, evaluates adult cohort survival with the Kaplan-Meier estimator and the log-rank test, and calculates sample size for lifespan studies using Schoenfeld’s formula. The quantitative components were checked against independent references, including R’s survival package and manual calculations. Ongoing work is focused on validating the calibrated model using cohorts specifically bred for this purpose. DrosoTracker runs entirely in the browser, stores data locally, and is available in English and Spanish.

## Introduction

In the daily routine of a laboratory working with the Drosophila melanogaster model, there are various tasks that must be carried out for the proper husbandry of the model^1^. The question "what day do I have to do this?" is a constant one, and the answer depends on rearing temperature, as well as on other conditions that are usually held constant, such as density^2^, food and humidity^3^. The same line completes development in different times at 18 °C and at 29 °C^4^. This thermal dependence makes Drosophila a highly tractable system, since development can be accelerated or slowed by varying a single parameter.

Estimating the different periods of the fly’s life cycle is an important detail because the time windows for some tasks are short^1^. A window that is missed forces the researcher to wait for the next cohort or to discard an experiment. Moreover, planning an experiment backwards from a target date requires an inverse calculation which is simple to do once, but it is repeated for every cross, genotype and temperature, and it is that repetition that opens the door to avoidable errors whose cost is lost time.

Tools such as drosophila.me^5^, LabDB^6^ or flyRoom^7^ currently serve as managers for keeping track of a laboratory’s stocks and crosses, planning crossing schemes and importing stocks from public centers such as FlyBase^8^ and the Bloomington Drosophila Stock Center (BDSC)^9^, among other functions. These address a real and central problem of fly work, namely the administration of a living collection and its traceability. Experimental work, on the other hand, poses a different problem. There, the goal is to obtain flies of a given genotype and age on a given date. Rearing temperature is chosen according to the experiment and is even deliberately modified during development, as occurs with thermosensitive expression systems, and each of those changes shifts all subsequent dates. A fixed interval cannot account for that dynamic. We did not find, however, a tool that predicts development from an explicit thermal model derived from published data, that allows that model to be corrected with a laboratory’s own observations, and that integrates the tracking of lifespan cohorts.

Here we present DrosoTracker, a tool that addresses this gap. At its core is a thermal-summation model whose constants we did not adopt from a table of rounded values, but rather recalibrated by regression from the data published by Powsner^10^. On top of that literature model, DrosoTracker learns two optional corrections from the user’s own observations. One captures the particular conditions of the laboratory through a wild-type reference strain, and the other, nested within the former, is specific to each genotype. A shrinkage estimator pulls back toward the literature model when there are few data and converges toward the observations when there are many. From that model, the application automatically generates the husbandry tasks corresponding to each cross (collecting virgins, clearing parents, transferring adults to fresh food), tracks the survival of adult cohorts with the Kaplan-Meier estimator and the log-rank test, and helps to size lifespan experiments before starting them. The novelty of DrosoTracker is that its developmental model is anchored in traceable primary data and calibrates itself to each user’s conditions, instead of assuming that a single pair of constants holds for any strain and laboratory.

## Methods

### Software architecture

DrosoTracker is a single self-contained HTML document, written in JavaScript and CSS without frameworks, with no build step and no runtime dependencies beyond web fonts. Data are stored in localStorage and do not leave the device. The application can be installed as a Progressive Web App (PWA), works offline through a service worker, and is available in English and Spanish.

Since an error in a calculation can go unnoticed, as it produces results that seem reasonable, we verify the code automatically. We include a set of 130 automated checks (release v1.0, archived at Zenodo, DOI: 10.5281/zenodo.21867404) that evaluate the calculation functions extracted directly from the released HTML file, so that the code under test is the code that ships. These checks compare results against values computed by hand and against independent references. They run every time we modify the code, so any change that affects a calculation is detected immediately. Additionally, the calibration analysis with literature data runs in a separate script, written in R. This script reproduces all the values and the figure we report from the primary data and checks that they match the published ones.

### Developmental model and calibration equation

We adopted the classical thermal-summation model for ectotherm development^11^. The linear formulation with a lower threshold and a thermal constant, along with its estimation by regression, is standard in the entomological literature^12–14^.

The model assumes that the time required to complete development from egg to adult is inversely proportional to how much temperature θ exceeds a lower threshold T_0_:

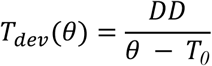

Where DD is the thermal constant, expressed in °C·days. The difference (θ - T_0_) indicates the effective degrees accumulated each day, and DD is the total amount of heat the individual needs to gather to complete development. In other words, the developmental rate is linear with respect to temperature.

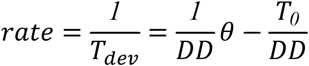

This reformulation is what allows both constants to be estimated by a simple linear regression of rate against temperature, since:

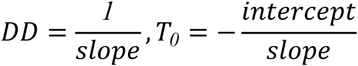

For the calibration, we fitted this line to the egg-to-adult developmental times measured by Powsner^10^. We calculated these times as the sum of the egg-larval period and the pupal period, averaging both sexes. We limited the fit to the linear range, roughly between 15 and 28 °C (n = 13), and excluded points above about 28 °C. These points are shown in Figure 1 but do not contribute to the fit because, at that high temperature, the relationship between rate and temperature changes. We obtained the confidence intervals for T_0_ and DD using the delta method from the covariance matrix of the fit. The outcome is T_0_ = 11.78 °C and DD = 116.38 °C·days (R² = 0.997). The total developmental time is split into stages based on the proportions Powsner measured at 25 °C. These are embryo 9.8 %, larval 44.8 %, and pupal 45.4 %. We rescale these figures with the predicted total duration. The larval period is further subdivided into L1:L2:L3 = 1:1:2. This division comes from the FlyBase developmental staging^8^, which is 25:23:48 hours at 25 °C rather than from Powsner, who measured the larval period as a single block. The boundaries of each stage at any temperature are calculated by rescaling these fixed proportions over T_dev(θ)_. Users see this information on a timeline that shows the predicted date of each transition (Figure 3A, B).

**Figure 1.**
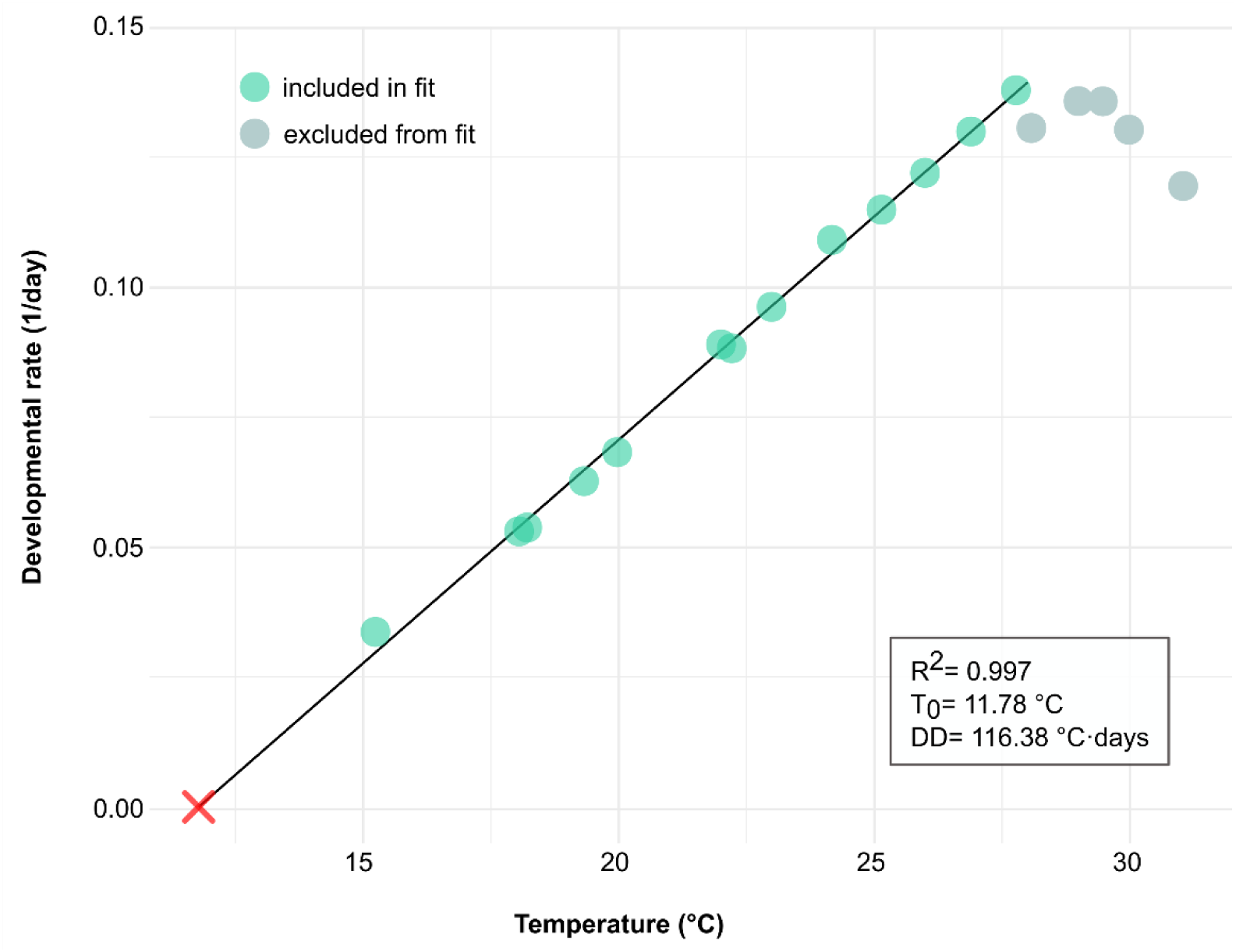
Figure 1. Calibration of the thermal-summation model from primary data. Developmental rate as a function of rearing temperature, calculated from the data of Powsner1. The green points are those included in the fit (n = 13, approximately 15-28 °C), and the grey ones represent those excluded, since above that range it departs from linearity and even reverses. The line corresponds to the linear regression fit, and the red cross marks its intersection with the x-axis, which defines the lower developmental threshold T₀. From the slope and the intercept, the constants of the base model are obtained (T₀ = 11.78 °C; DD = 116.38 °C·days; R² = 0.997).

### Two-level calibration: laboratory and genotype

Deviations from the literature model have two distinct sources. First, the rearing conditions in a specific laboratory affect all strains (Figure 2). Second, the genotype influences a particular line. DrosoTracker distinguishes between these sources through an optional two-level calibration. The user can first select a wild-type reference strain and calibrate the model with it. The factor ƒ_lab_ becomes the adjusted baseline for the laboratory and applies to any strain that lacks its own calibration. Next, each genotype is calibrated against that baseline. The estimator for a genotype moves toward ƒ_lab_ instead of the literature model (ƒ = 1). This way, a strain with limited data is adjusted to the user’s conditions and not to Powsner’s.

**Figure 2.**
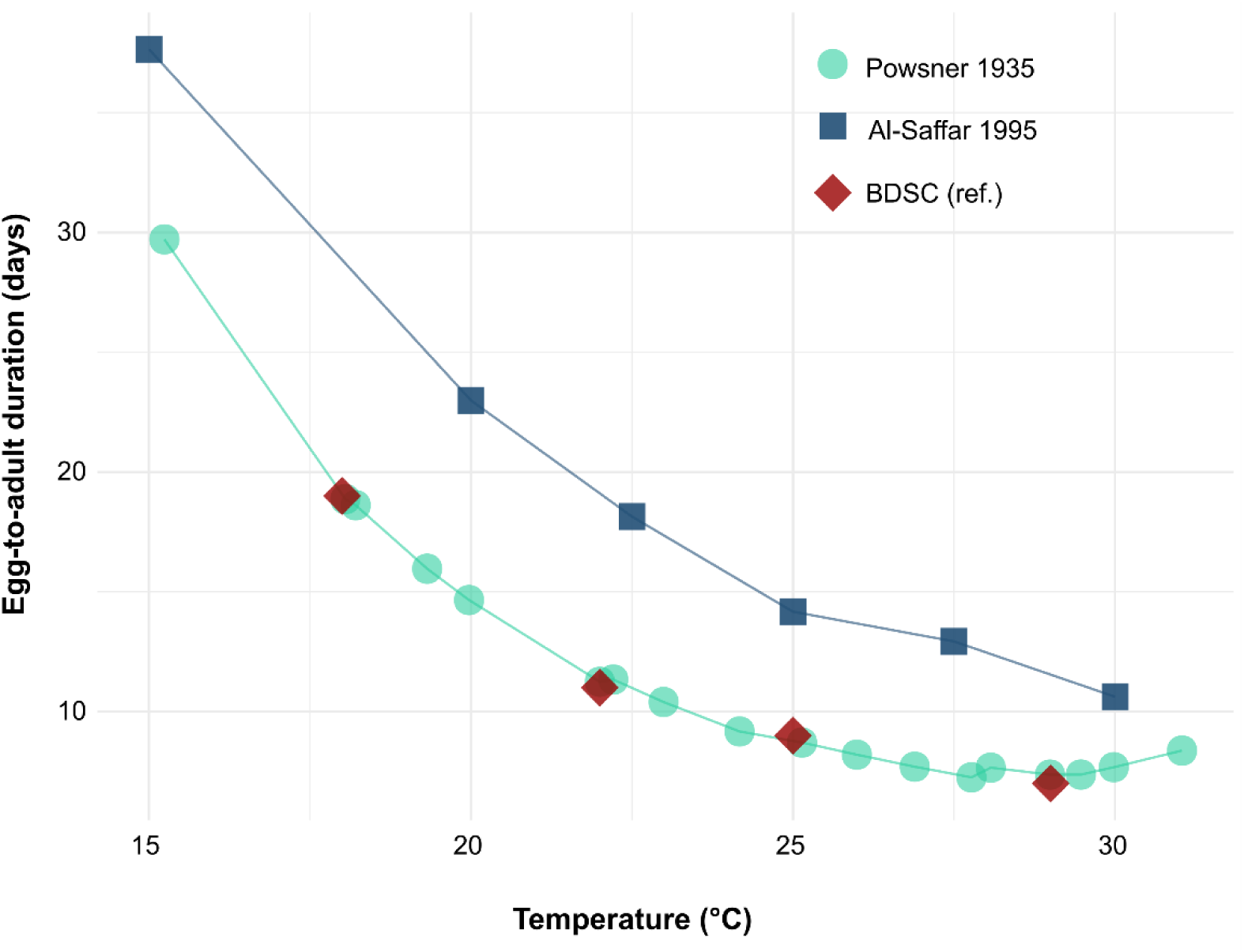
Variation between studies in egg-to-adult developmental time. Duration of egg-to-adult development as a function of temperature, according to three sources: Powsner10, which we used as the basis for the calibration; Al-Saffar25; and the reference values of the BDSC9. The values of Al-Saffar are systematically longer than Powsner’s, on the order of 60 to 70 % between 20 and 27.5 °C. The BDSC reference values, in contrast, closely match those of Powsner across the whole range. The comparison illustrates that thermal constants depend on rearing conditions as much as on genotype and provides the basis for the user-level calibration that DrosoTracker implements.

**Figure 3.**
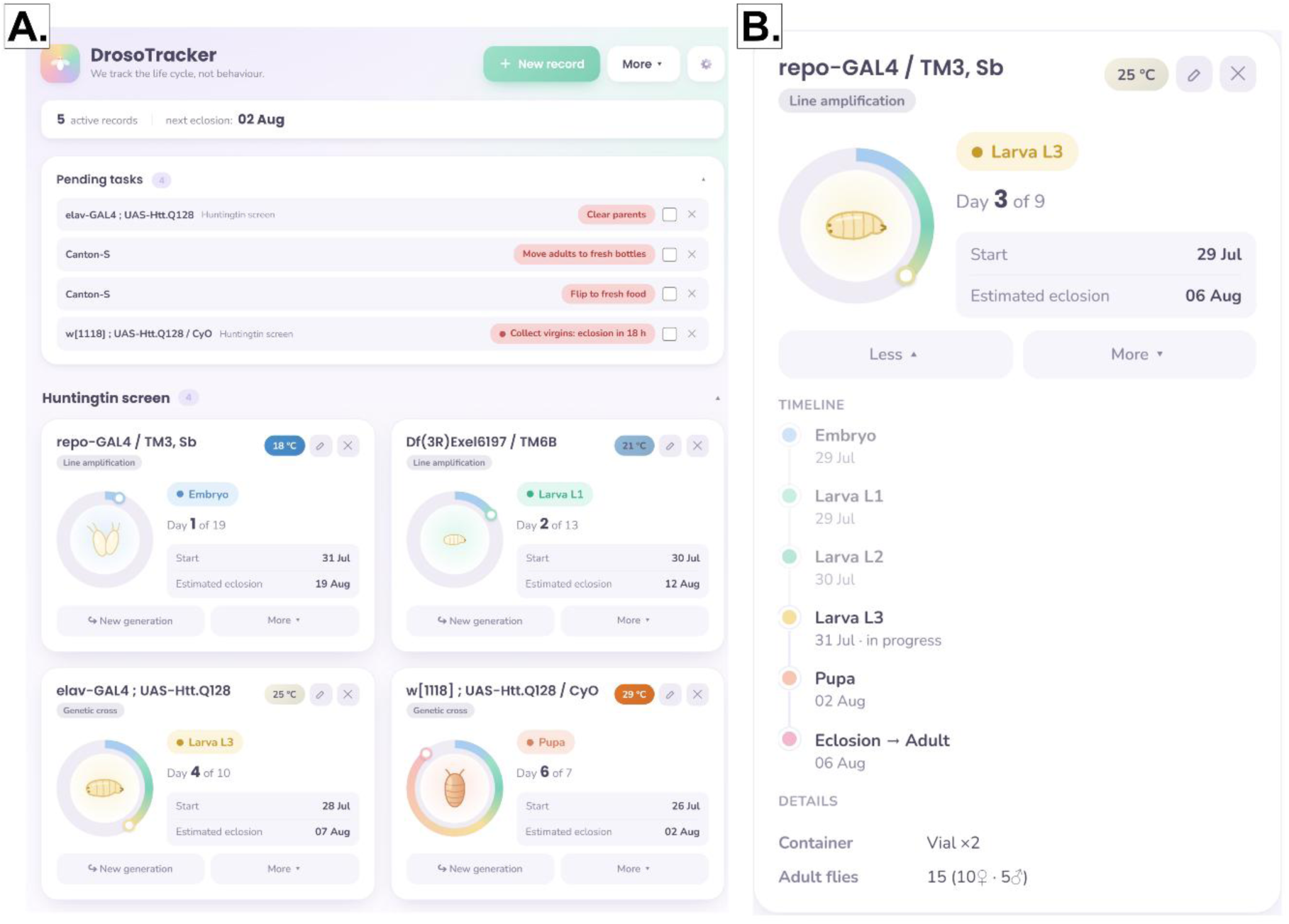
Model-derived husbandry scheduling. (A) Main view of the application. Active records are grouped by project, and each card reports the genotype, the rearing temperature, the record type, the current developmental stage, the day within the predicted total duration and the estimated eclosion date. (B) Expanded view of a single record, showing the timeline of predicted stage transitions with their dates, from embryo through the three larval instars and the pupal period to eclosion, together with the rearing temperature and the recorded contents of the vial. All data shown are for demonstration purposes.

At both levels, this tool scales DD with a single multiplicative factor ƒ, while keeping T_0_ fixed, since it cannot be reliably estimated from observations at a single rearing temperature. Learning only one parameter prevents overfitting with few observations and allows the correction to apply to all temperatures. The shape of the thermal curve remains the same; only its magnitude changes. Each observation gives a starting point for the egg-lay (t_0_) and the time until half of the cohort ecloses (t_50_). From this, an observed factor is derived:

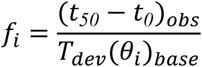

To combine the observations of a given level into a single factor, we use a random-effects estimator based on the literature model. This estimator adjusts to the user’s data as these accumulate and includes four steps. First, each observation has an associated uncertainty σ that depends on how eclosion was recorded, whether by counting or visual estimation, and on how well t_0_ is known. Second, the batch, essentially the cross, constitutes the experimental unit. We combine observations within the same replicate using inverse variance, with a minimum limit on the variance between them. Third, we check whether the batches agree with each other beyond their own measurement error. We measure this variation using Cochran’s Q^15^ and estimate the between-batch variance by the method of DerSimonian and Laird^16^, also reporting it as I²^17^. When variation is high, the confidence interval widens, and the genotype is marked as inconsistent. Fourth, the combined estimate moves toward the corresponding baseline through a normal distribution centered on it, which is ƒ_lab_ when there is a calibrated reference strain and ƒ = 1 when there is not. This method helps keep sparse data close to the baseline while allowing precise data to converge toward the observations.

The software displays the status of each genotype (uncalibrated, calibrating, calibrated from three independent batches, or inconsistent) together with its 95 % interval, and warns when all observations come from a single batch, a situation in which the effect of the genotype is confounded with that of the batch (Figure 4A).

**Figure 4.**
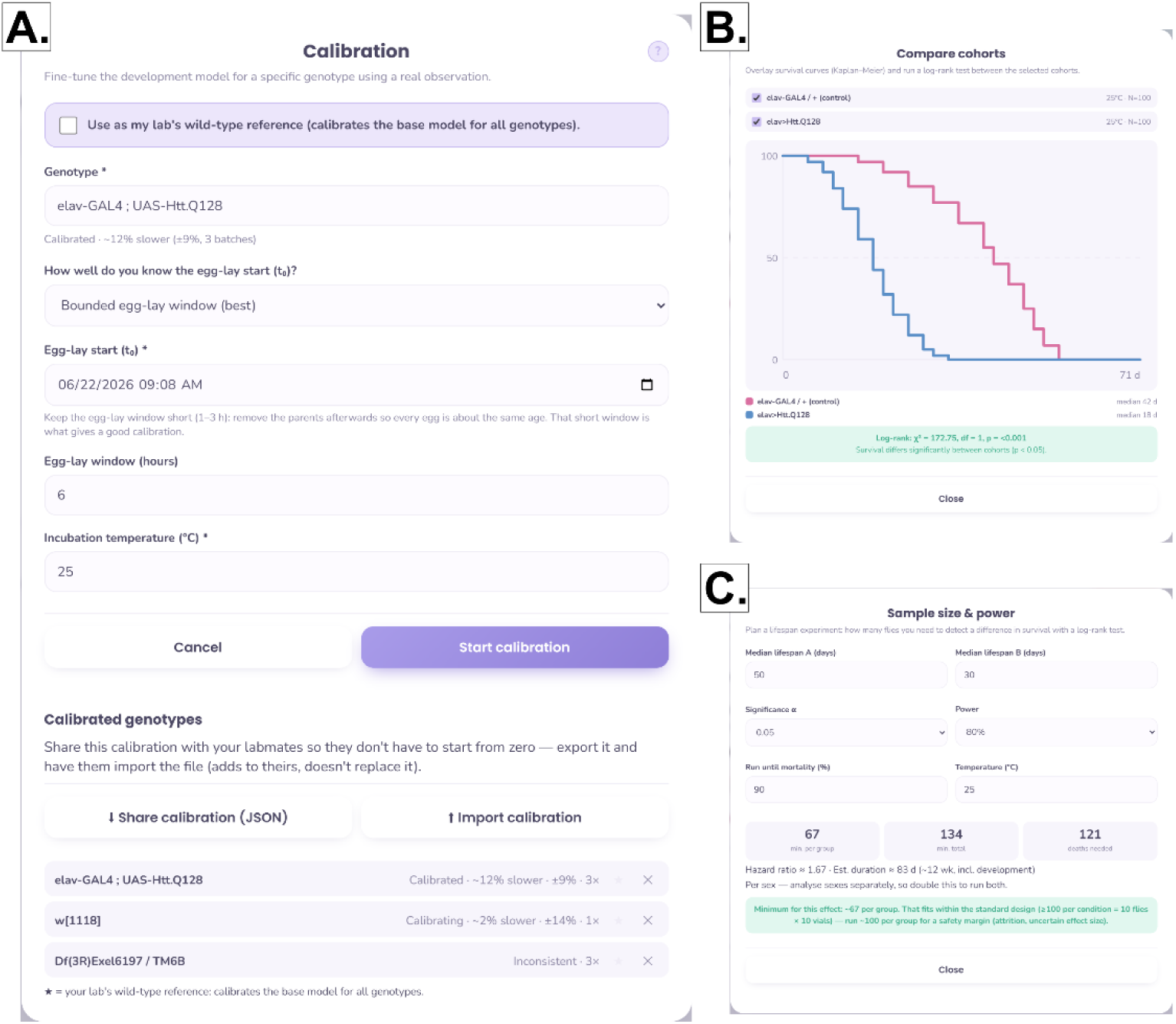
Genotype calibration and lifespan modules. (A) Calibration form and status of the calibrated genotypes. (B) Cohort comparison. Kaplan-Meier curves for the selected cohorts are overlaid, with the median of each group and the result of the log-rank test. (C) Sample-size and power calculator. All values shown are illustrative.

### Survival analysis

Adult survival is estimated using the Kaplan-Meier estimator^18^:

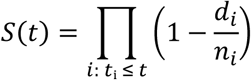

Here dᵢ are the deaths observed at event time tᵢ and nᵢ the number of individuals at risk immediately before tᵢ. The product runs over all event times up to and including t. Flies that escape or die accidentally during food transfers are censored, meaning they do not count as events but still provide information until they are lost. The application reports median and maximum survival and allows for group data export for analysis.

Cohorts are compared using the log-rank test introduced by Mantel^19^. For G groups, the statistic is a quadratic form over a reduced system of G - 1 dimensions^20,21^.

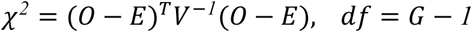

O and E are the vectors of observed and expected deaths for any G - 1 of the G groups, and V is the corresponding (G - 1) × (G - 1) submatrix of the multivariate hypergeometric variance-covariance matrix. One group is dropped because the full G × G matrix is singular and cannot be inverted. The expression reduces to the familiar two-group formula, and the p-value is calculated using the regularized upper incomplete gamma function (Figure 4B).

### Experiment planning

The size of lifespan experiments is calculated with Schoenfeld’s event formula^22^ for the log-rank test, assuming 1:1 allocation between groups:

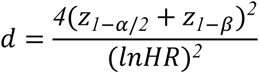

Where d is the total number of events (deaths) required. The effect size is based on two median lifespans, which helps derive the hazard ratio (HR). This ratio compares the instantaneous risks of death between the two groups. An HR of 2 means one group has double the risk of dying compared to the other. The calculation uses an exponential model (HR = m_1_/m_2_), which implies a constant risk of death as age increases. In *Drosophila*, this does not hold strictly, since mortality follows a Gompertz-type curve^23^. This formula is for planning purposes and should not be interpreted as a definitive result.

Events are converted into the number of flies per group based on the expected fraction of events. The estimated duration of the experiment adds the egg-to-adult developmental time to the time needed for the cohort to reach a target mortality fraction q, which the user sets in the interface (default 90 %). That time is projected from the median lifespan m using the quantile function of a Weibull survival model with shape parameter k:

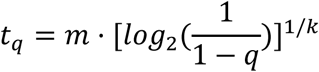

An exponential model (k = 1) is not appropriate here, because in Drosophila mortality accelerates with age rather than remaining constant^23^, and a Weibull with k > 1 reproduces that compressed tail while remaining analytically simple^24^. We fixed k = 3, which is equivalent to assuming that 90 % mortality is reached at approximately 1.5 times the median lifespan, as opposed to 3.3 times the median under an exponential model. The choice of k therefore has little influence on planning decisions, while the departure from the exponential model does. As with the sample-size calculation, this estimate is for planning purposes only (Figure 4C).

### Verification

We checked each quantitative component against an independent reference. The Kaplan-Meier estimator was verified against R’s survival:survfit, on a censored dataset with a known median. The log-rank statistic was compared with a worked example calculated by hand with interleaved events and with known critical values of χ². The sample-size formula was checked against the standard result of 66 events for an HR = 2, with α = 0.05 two-sided and 80 % power. Finally, the calibration analysis automatically verifies that the reported values match the published ones.

## Results

We first fitted the thermal-summation model to Powsner’s data. Over the linear part of the response, developmental rate increased with rearing temperature along a straight line (n = 13, approximately 15 to 28 °C), and the slope and intercept of that line define the two constants of the base model. The fit yields a lower developmental threshold of 11.78 °C (95 % CI 11.39–12.18) and a thermal constant of 116.38 °C·days (95 % CI 112.20–120.56), with R² = 0.997 (Figure 1). Points above 28 °C were excluded from the fit (plotted in grey) because the rate does not continue to rise with temperature and eventually decreases. These constants predict an egg-to-adult duration of about 8.8 days at 25 °C, 12.6 days at 21 °C, 18.7 days at 18 °C and 6.8 days at 29 °C. Rescaling those totals based on the stage proportions Powsner measured at 25 °C (embryo 9.8 %, larval 44.8 %, pupal 45.4 %) places the larval-pupal transition at about day 4.8 at 25 °C, which is the type of date a scheduling decision relies on.

Before building any scheduling logic on a single pair of constants, we asked how far those constants can be expected to travel. Comparing three independent sources of egg-to-adult times across the same thermal range (Figure 2), the values reported by Al-Saffar^25^ are systematically longer than Powsner’s, by roughly 60 to 70 % between 20 and 27.5 °C, whereas the reference values of the Bloomington Drosophila Stock Center track Powsner closely throughout. A discrepancy of that size between two published datasets is larger than most of the effects a scheduling tool is expected to resolve. We take this as the empirical justification for the two-level calibration described in Methods, in which a wild-type reference strain absorbs what is specific to a laboratory and each genotype is then estimated relative to that baseline rather than relative to Powsner’s original strain.

The application turns the resulting model into a working schedule (Figure 3). Each record stores a genotype, a rearing temperature and a record type (e.g., a genetic cross or a line amplification), and from these inputs it provides the current developmental stage, the day within the predicted total duration and the estimated eclosion date. Records held at different temperatures run side by side, and each receives its own timeline. In the demonstration data shown, records at 18, 21, 25 and 29 °C have predicted durations of 19, 13, 10 and 7 days, respectively (Figure 3A). The 25 °C record shows how calibration affects the schedule, since it belongs to a genotype estimated to be about 12 % slower than the baseline and is therefore given 10 days rather than approximately 9 days that the uncalibrated model provides for a genotype without its own calibration. Most pending tasks are not entered by the user; they come from these predictions. Virgin collection is announced with the eclosion window still open, parents are cleared once the larvae are established, and adults are flipped to fresh food on schedule. Changing the rearing temperature of a record shifts every subsequent date and every associated task without further intervention. Finally, expanding a record shows the complete sequence of predicted transitions with their dates, from embryo through the three larval instars and the pupal period to eclosion, along with the recorded contents of the vial (Figure 3B).

Calibration is recorded as an observation instead of a parameter (Figure 4A). The user provides the length of the egg-lay window and the incubation temperature. They can also mark the line as the laboratory’s wild-type reference, in which case it recalibrates the baseline for all other genotypes. Each genotype’s status is shown along with the direction and magnitude of its deviation, its 95 % interval and the number of independent batches supporting it. The three states the estimator can achieve are visible in the demonstration data: A genotype with three matching batches is labelled as calibrated and reported as approximately 12 % slower (± 9 %); a genotype with a single batch remains in a calibrating state with a visibly wider interval (± 14 %); and a genotype whose batches disagree beyond their own measurement error is marked as inconsistent even though three batches are available. This is the outcome we aimed for because, in this case, the point estimate is not what the user should rely on. Calibrations can be exported and imported as a JSON file, so that a laboratory-level calibration needs to be set up only once.

The lifespan module covers the two points at which a survival experiment needs quantitative support: comparing cohorts that have already been run, and sizing one that has not (Figure 4B, C). Cohorts are overlaid as Kaplan-Meier curves and compared with the log-rank test, with medians reported for each group and flies lost during transfers censored rather than counted as deaths. In the sample cohorts, two groups of 100 flies with medians of 42 and 18 days give χ^2^ = 172.75 with 1 degree of freedom and p < 0.001. These numbers come from demonstration data and illustrate what the interface reports.

The planning calculator runs the same logic backward. In the example, two hypothetical median lifespans of 50 and 30 days give a hazard ratio of about 1.67 and, at α = 0.05 and 80 % power, require 121 events. Assuming the cohort is followed until 90 % mortality, the default in the interface, the tool converts that into a minimum of 67 flies per group and an estimated duration of about 83 days at 25 °C, egg-to-adult development included. It also indicates whether that requirement fits the field standard of roughly 100 flies per condition, and reminds the user that sexes are analyzed separately, so the totals double if both are to be run.

A wrong date looks exactly like a right one, so we verified each quantitative component against an independent reference. All 130 automated checks pass, and continuous integration re-runs them on every commit. The Kaplan-Meier implementation reproduces R’s survival:survfit exactly, the log-rank statistic matches the hand-calculated example, and the sample-size routine returns the expected 66 events. Separately, an R script regenerates the constants and Figure 1 from Powsner’s data and verifies that the reported values match the published ones, so the calibration shown here is reproducible from its source rather than transcribed.

What these results establish is that the model is correctly fitted, correctly implemented and correctly propagated into the schedule; they do not yet establish how accurate its predictions are in practice. That evaluation is currently under way and is not included in this version. We are rearing cohorts across the linear range of the model, in batches independent of those used for calibration, and will contrast predicted with observed eclosion times under the uncalibrated literature model and under the calibrated one, so that the improvement attributable to calibration can be quantified rather than assumed. Those data will be reported as an additional figure in a subsequent version of this manuscript.

## Discussion

This study describes DrosoTracker, a browser-based tool for scheduling Drosophila melanogaster husbandry tasks and planning experiments, whose developmental model was derived from published data^10^ and can be adapted to the conditions of each laboratory. Existing laboratory management tools such as drosophila.me^5^, LabDB^6^ and flyRoom^7^ address the logistics of maintaining a fly collection: they track stocks and crosses, help design crossing schemes, and import stock information from public centers. None of them, however, predicts when a given cross will reach a target developmental stage, or learns from a laboratory’s own rearing conditions. DrosoTracker does not replace these tools: it does not track pedigrees, import stocks from public stock centers, or manage a laboratory’s full collection, and a user working across both kinds of problems will still need a stock manager alongside it. It instead focuses narrowly on the developmental-timing problem that stock-management tools were not designed to solve, so the two kinds of software are complementary rather than competing; a natural extension would be for a stock database to import DrosoTracker’s calibrated schedules, or vice versa. Within that narrower scope, DrosoTracker’s husbandry tasks are not entered manually, as they would be in a virtual calendar, but derived from the thermal-summation model and adjusted to the rearing temperature. In addition, its lifespan module, detailed in Methods, handles cohort survival analysis and experiment sizing. All of this takes place within an open-source platform, with no registration or installation, and the data remains on the user’s device. This design has a corresponding cost: because no data leave the device, sharing a calibration among the members of a laboratory currently requires manually exporting and importing a JSON file (Figure 4A) rather than syncing automatically, a trade-off we judged acceptable for privacy and simplicity but one that could be revisited if multi-user support becomes a priority.

DrosoTracker predictions are designed to guide day-to-day laboratory decisions. Therefore, in addition to having a biologically grounded model, it is essential that users have confidence that the software correctly implements the calculations. To verify this aspect, we performed 130 automated checks using hand calculations and independently obtained values. We also developed an R script that reproduces the calibration analysis and the corresponding figure directly from the original data. Together, these verifications reduce the likelihood that calculation errors go unnoticed. The accuracy of the calculations performed by the software, however, should be considered independently from the biological validity of the thermal summation model, which is analyzed in the following section.

Using primary data, we fitted the thermal-summation model^11^ by linear regression to the egg-to-adult developmental times measured by Powsner, thereby obtaining the constants of the base model (T_0_ = 11.78 °C, DD = 116.38 °C·days). As Sewall Wright noted in Powsner’s own paper^10^, the values obtained hold for the particular strain Powsner worked with and cannot be carried over to other stocks. In addition, a second source of variation is the rearing conditions (e.g. food^26^, humidity^3^, incubator accuracy, handling, and population density^2^). For this reason, DrosoTracker allows a wild-type line to be used to calibrate the conditions of each laboratory, which are inherent to all the lines of its stock, and each genotype can then be calibrated relative to that baseline rather than to the literature model.

Among the parameters obtained by fitting the data, we find T_0_, which is the point at which the line of developmental rate against temperature crosses the x-axis. It does not represent a real physiological temperature below which development stops^27^. On the contrary, Powsner showed that development still occurs below this temperature^10^. He also noted that the relationship between rate and temperature is sigmoid, so that linearity is a valid approximation in the mid-range (15–28 °C)^1,25^. Near the upper end, developmental time stops decreasing and begins to increase, so that the model underestimates duration at the warm extreme. At 29 °C, a rearing temperature frequently used in fly work; the prediction falls approximately 15 hours below what Powsner observed^1^, and the error grows rapidly above that temperature. Fluctuating temperatures also modify developmental times. Under alternating temperature regimes, development proceeds faster than predicted. This phenomenon, known as the Kaufmann effect^25,28^, is a known and bounded bias that laboratory-level calibration absorbs empirically.

These thermal-range limitations are compounded by further assumptions the model makes beyond temperature itself. It assumes that the proportion of total development taken up by each stage remains fixed at the values measured at 25 °C (embryo 9.8 %, larval 44.8%, and pupal 45.4 %) and is simply rescaled with the predicted total duration. Although the larval period is subdivided into instars according to the proportion L1:L2:L3 = 1:1:2, taken from the FlyBase developmental staging^8^ (L1:L2:L3 = 25:23:48 h at 25 °C), this subdivision is not a parameter fitted by our model, and it is known to depend on rearing conditions. Individually reared larvae followed by automated tracking on a yeast-supplemented agarose medium show a proportionally longer third instar, with molts at 1.9 and 3.5 days and pupariation at 7.8 days after hatching, equivalent to L1:L2:L3 = 1:0.8:2.3^29^. Finally, the model treats temperature as constant and ignores larval density^2^, humidity^3^ and nutritional quality^26^, factors that measurably shift developmental time. Predictions should therefore be interpreted as calibrated expectations that structure a schedule, not as substitutes for observation. Lastly, the pending task of this work is the empirical validation of the model, that is, contrasting predicted eclosion times with those observed in our own cohorts reared at different temperatures and in batches independent of those used for calibration.

None of the individual components behind DrosoTracker, the thermal-summation model, the Kaplan-Meier estimator, or Schoenfeld’s formula, is new; the contribution lies in calibrating and integrating them into a single, self-calibrating scheduling tool. It is worth noting, in turn, that this integrative approach is not exclusive to the fruit fly^30^. The thermal-summation model describes the development of any ectotherm reared at controlled temperatures, so it can be applied to other species with their own constants. In practice, DrosoTracker is best suited to laboratories that keep eclosion records for at least a few batches of a wild-type strain, since this is what allows the laboratory- and genotype-level calibrations to converge away from the literature baseline; without such records the tool still generates a schedule, but one that reflects Powsner’s original strain rather than the user’s own colony. All of this makes DrosoTracker a useful tool for day-to-day work with Drosophila melanogaster and a basis for extending the same approach to other model organisms.

## Declaration of generative AI use

The released software contains no artificial-intelligence components: DrosoTracker is fully deterministic, built from a published model and the user’s own observations, with no language model or external service involved in any of its calculations. During development, an AI coding assistant (Anthropic Claude, via Claude Code) was used under the authors’ direction to assist with the software implementation, the analysis and test scripts, and the language editing and translation of the manuscript. The authors directed the work, verified the quantitative results against the independent references described in Methods, and take full responsibility for the entire content.

## Data availability statement

The DrosoTracker application, the calibration script, the estimator simulation and sensitivity scripts, the test suite and the primary data used to fit the thermal-summation model are openly available at https://github.com/gersonasti/Drosotracker under the MIT licence, which permits use, modification and redistribution provided that the copyright notice and the licence text are retained. The release described here (v1.0) is archived at Zenodo, DOI: 10.5281/zenodo.21867404. The live application runs at https://drosotracker.maimonides.edu. No restrictions apply to data access.

## CRediT author contributions

**Gerson Smith Asti Tello:** Conceptualization, Methodology, Software, Formal analysis, Investigation, Data curation, Visualization, Writing – original draft.

**Mariana Melani:** Validation, Writing – review & editing, Supervision.

**Ana Clara Liberman:** Validation, Writing – review & editing, Supervision, Funding acquisition, Project administration.

